# Voxel-wise tracer kinetic model selection for DCE-MRI measurements of blood-brain barrier leakage

**DOI:** 10.64898/2026.05.29.728495

**Authors:** Olivia A. Jones, Ben R. Dickie, Michael Berks, Sarah Al-Bachari, Hedley C. A. Emsley, Laura M. Parkes

## Abstract

**Purpose:** To apply voxel-wise tracer kinetic model selection, characterise the spatial distribution of best-fitting models across the brain, and evaluate whether model selection improves sensitivity for differentiating normal-appearing tissue from pathological tissue compared to the Patlak model.

**Methods:** Extended Tofts, Patlak, and intravascular models were fit to DCE-MRI data from stroke survivors and controls, as well as simulated data. The best-fitting model was chosen for each voxel using the Akaike Information Criterion, and model selection *K*^trans^ (estimates from the best-fitting model for each voxel) compared to Patlak model *K*^trans^.

**Results:** In simulated data, the Extended Tofts model was best-fitting at *K*^trans^>10^−3^ min^-1^, where the Patlak model systematically underestimated *K*^trans^. Patlak was optimal at *K*^trans^ between 10^−4^-10^−3^ min^-1^, where Extended Tofts estimates had greater variability. The intravascular model was selected for *K*^trans^∼10^−4^ min^-1^. The Patlak model was chosen in most control voxels. In chronic stroke, the Extended Tofts model was preferred in most cortical and white matter hyperintensity voxels, while the Patlak model was selected in most deep grey matter and normal-appearing white matter voxels. Model selection *K*^trans^ estimates were significantly greater than Patlak estimates in the cortex and white matter hyperintensities, with greater inter-patient variability, likely reflecting biological variability in blood-brain barrier leakage resulting from stroke.

**Conclusion:** Voxel-wise model selection may provide more accurate estimates of a wider range of *K*^trans^ values than any single model, revealing greater differences between normal and pathological tissue and offering a more sensitive and physiologically appropriate framework for DCE-MRI analysis of blood-brain barrier dysfunction.

## 1 Introduction

Blood-brain barrier (BBB) dysfunction is a hallmark of neurological disease and an increasingly important topic of research. *In-vivo* study of the BBB in humans is commonly performed using dynamic contrast-enhanced MRI (DCE-MRI), where a paramagnetic contrast agent is administered part-way through a dynamic image series. Mathematical modelling of the contrast agent concentration over time enables the estimation of physiological parameters relating to BBB integrity, such as the contrast agent volume transfer constant, *K*^trans^. It is becoming more popular to perform this modelling on a voxel-wise basis to produce parametric maps of BBB leakage; voxel-wise analysis has been employed across a wide range of pathologies including intracerebral haemorrhage,^1^ ischaemic stroke,^2,3^ Parkinson’s disease,^4^ traumatic brain injury,^5,6^ and small vessel disease.^7,8^

There are several possible models to choose from,^9^ as shown in Figure 1, and these models are nested based on a series of simplifying assumptions: the Extended Tofts model^10^ models bi-directional tracer exchange between finite intravascular and extravascular spaces, with the assumption that the tissue or voxel is well perfused. This simplifies to the Patlak model^11^ if there is no clearance or back-flux of the tracer after it has left the intravascular compartment, and simplifies further to an ‘intravascular’ model^12^ in the case where there is no tracer leakage, valid for a completely intact BBB. It is conventional to fit a single model to the entire brain, and the Patlak model in particular has been shown to fit well in cases of subtle leakage^13^ and is widely applied.

**Figure 1.**
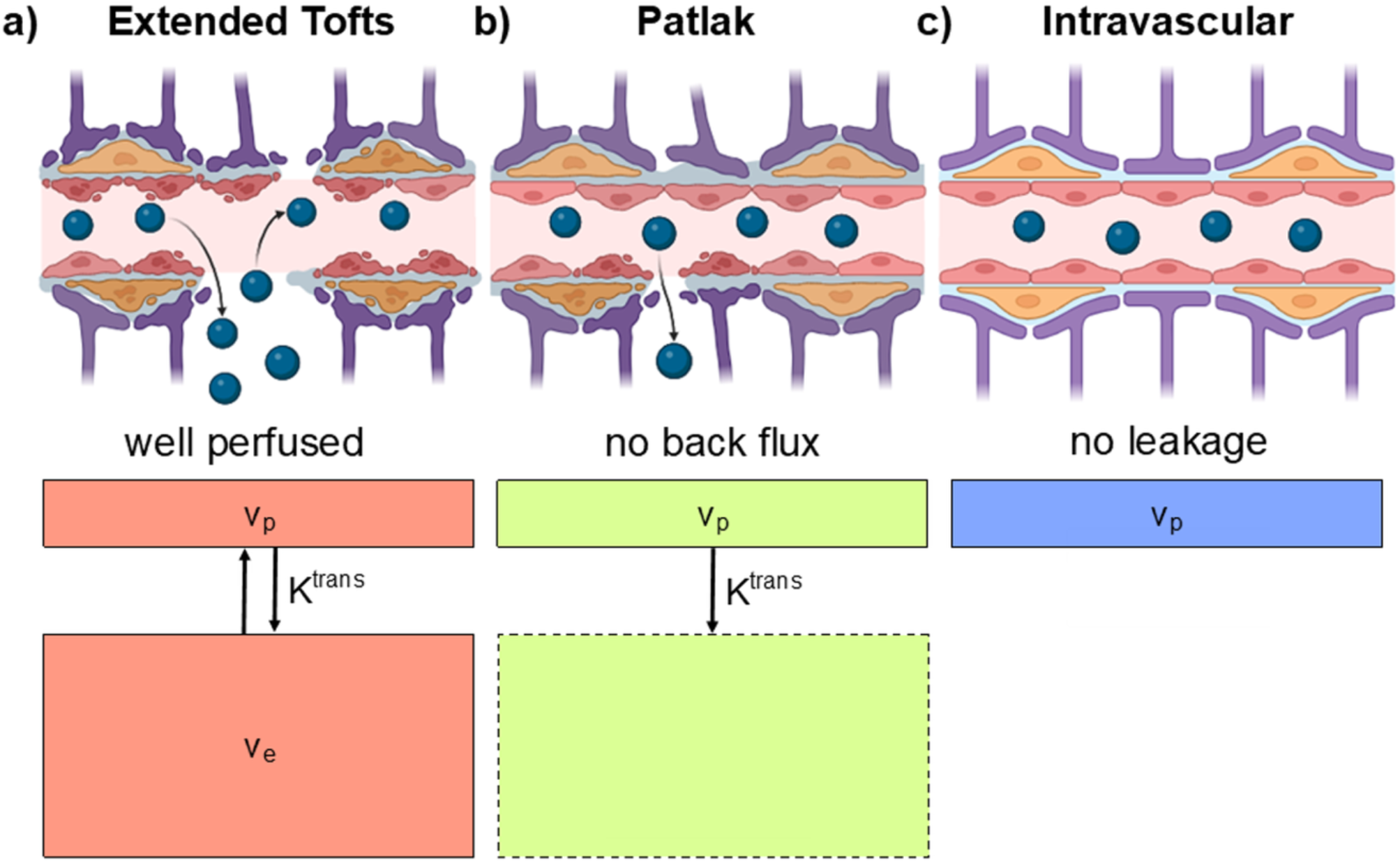
DCE-MRI Tracer Kinetic Models. Diagram representing the nested tracer kinetic models used in analysis and their associated assumptions. **(a)** The full Extended Tofts model, which models exchange of a tracer between the blood plasma compartment and a finite interstitial space in a well perfused voxel. **(b)** A Patlak model of tracer leakage, which assumes no backflux/clearance from the interstitial space into the blood plasma compartment. **(c)** An intravascular model, which assumes no tracer leakage.

The choice of model directly impacts estimates of leakage parameters and consequently the interpretation of results. The accuracy of model assumptions is driven by the voxel/tissue pathophysiology. BBB permeability is unlikely to be homogenous across the brain,^14^ and this is particularly true in aging or neurological disease, where disruption may be severe at a lesion site while remaining subtle in tissue that appears otherwise normal. The spatial extent and severity of BBB dysfunction cannot be known in advance of the model fitting, this is precisely what the experiment aims to measure. Selecting a single model for a whole brain or tissue region therefore risks errors and mischaracterisation of voxels where the underlying model assumptions do not hold. Voxel-wise selection of the best-fitting model may overcome this problem, with the greatest potential benefit in cases where focal injury/lesions coexist with diffuse, subtle damage, such as stroke. Beyond improving the accuracy of BBB leakage estimates, the preferred model itself may reveal useful information, as tissue properties could be inferred from the validity of model assumptions, for example the intravascular model fitting best implying an intact BBB.

Here, we have applied voxel-wise model selection to simulated and in-vivo DCE-MRI data from stroke patients and healthy controls with the following aims:

1. Evaluate how vessel permeability and noise influence model selection and *K*^trans^ estimates in simulated DCE-MRI data.
2. Characterise the spatial distribution of best-fitting models across the cortex, deep grey matter (DGM), normal-appearing white matter (NAWM), and white matter hyperintensities (WMH) in stroke survivors and controls.
3. Compare *K*^trans^ estimates derived from voxel-wise model selection to those from conventional Patlak fitting.
4. Assess whether voxel-wise model selection better distinguishes pathological from normal-appearing tissue compared to conventional Patlak fitting. We hypothesise that differences will be greater using the model selection approach than Patlak fitting alone.

## 2 Materials and methods

### 2.1 Quantities, Processes and Model Definitions

All quantities, processes and model definitions are OSIPI CAPLEX compliant.^15^ CAPLEX definitions can be accessed by clicking on quantity, process or model hyperlinks.

### 2.2 Simulations

The first aim of this study was to evaluate how vessel permeability and noise influence model selection and *K*^trans^ estimates in simulated DCE-MRI data. To address aim 1, a synthetic signal time-course, *S*_t_(*t*), was produced using the [two-compartment exchange model](https://osipi.github.io/OSIPI_CAPLEX/perfusionModels/#2CXM),^12^ which models bi-directional exchange between well mixed intravascular and extravascular compartments as well as a finite blood plasma flow, [*F*_p_](https://osipi.github.io/OSIPI_CAPLEX/quantities/#Fp), where the contrast agent [volume transfer constant across the BBB, *K*^trans^](https://osipi.github.io/OSIPI_CAPLEX/quantities/#Ktrans), is related to both [*F*_p_](https://osipi.github.io/OSIPI_CAPLEX/quantities/#Fp) and vascular permeability, [*PS*](https://osipi.github.io/OSIPI_CAPLEX/quantities/#PS), by 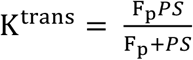^16^. Thus, this model has four free parameters: [*F*_p_](https://osipi.github.io/OSIPI_CAPLEX/quantities/#Fp), [*PS*](https://osipi.github.io/OSIPI_CAPLEX/quantities/#PS), the [fractional blood plasma volume,*v*_p_](https://osipi.github.io/OSIPI_CAPLEX/quantities/#v), and the [fractional extravascular extracellular volume,v_e_](https://osipi.github.io/OSIPI_CAPLEX/quantities/#v). Synthetic [indicator concentration time series,*C*_t_(*t*)](https://osipi.github.io/OSIPI_CAPLEX/quantities/#C), were generated by convolving a group-averaged vascular input function from a control group (described below) with the [two-compartment exchange model](https://osipi.github.io/OSIPI_CAPLEX/perfusionModels/#2CXM) using Madym’s^17^ madym_DCE_lite function in Python with the following input parameters: [*F*_p_](https://osipi.github.io/OSIPI_CAPLEX/quantities/#Fp) = 0.4 min^-1^, [*v*_p_](https://osipi.github.io/OSIPI_CAPLEX/quantities/#v) = 0.02, [*v*_e_](https://osipi.github.io/OSIPI_CAPLEX/quantities/#v) = 0.2, *T*_10_ = 0 s, and nineteen values for [*K*^trans^](https://osipi.github.io/OSIPI_CAPLEX/quantities/#Ktrans) = (1, 2, 3, 4, 5, 6, 7, 8, 9) x 10^−4^ min^-1^, and (1, 2, 3, 4, 5, 6, 7, 8, 9, 10) x 10^−3^ min^-1^. These parameters were chosen to approximately emulate the white matter in subtle (10^−4^ – 10^−3^ min^-1^) and moderate (10^−3^ – 10^−2^ min^-1^) BBB leakage regimes.^18,19^ The simulations were also repeated independently varying [*K*^trans^](https://osipi.github.io/OSIPI_CAPLEX/quantities/#Ktrans) with [*F*_p_](https://osipi.github.io/OSIPI_CAPLEX/quantities/#Fp), [*v*_p_](https://osipi.github.io/OSIPI_CAPLEX/quantities/#v), [*v*_e_](https://osipi.github.io/OSIPI_CAPLEX/quantities/#v), *T*_10_, and as a Monte Carlo simulation where all parameters were varied simultaneously, described in detail in the Supplementary Materials. The synthetic *C*_t_(*t*) was converted back to a [signal time-course,*S*_t_(*t*)](https://osipi.github.io/OSIPI_CAPLEX/quantities/#S), using Madym’s^17^ tissue_concentration function in Python with the same parameters as the clinical scan (described below in section 2.3.2) and an assumed [pre-contrast (native) longitudinal relaxation rate, *R*_10_](https://osipi.github.io/OSIPI_CAPLEX/quantities/#R10) of 1 s.

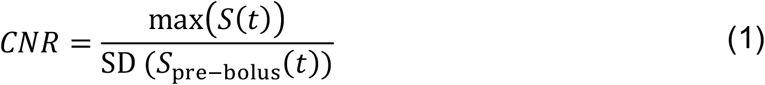

A range of random Rician noise was added to the simulated [signal time-series](https://osipi.github.io/OSIPI_CAPLEX/quantities/#S) using Madym’s^17^ add_rician_noise function in Python with standard deviations between 1 and 4. Rician noise was selected as it is generally regarded as more appropriate than Gaussian noise for MRI data,^20^ especially data with low SNR such as fast dynamic imaging.

The temporal [contrast to noise ratio (CNR)](https://osipi.github.io/OSIPI_CAPLEX/generalPurposeProcesses/#CalcCNR) was calculated for each time-series as the ratio between the maximum signal after contrast injection and the standard deviation of the signal in the 7 pre-bolus time points, described by equation 1. The noisy synthetic [signal time-series](https://osipi.github.io/OSIPI_CAPLEX/quantities/#S) were analysed using the same method as for the clinical data, as described below to produce estimates of [*K*^trans^](https://osipi.github.io/OSIPI_CAPLEX/quantities/#Ktrans) for the [Extended Tofts model](https://osipi.github.io/OSIPI_CAPLEX/perfusionModels/#ETM), [Patlak model](https://osipi.github.io/OSIPI_CAPLEX/perfusionModels/#Patlak), intravascular model, and model-selection method. This process was repeated 1000 times and the [median values](https://osipi.github.io/OSIPI_CAPLEX/generalPurposeProcesses/#CalcMedian) of noisy [*K*^trans^](https://osipi.github.io/OSIPI_CAPLEX/quantities/#Ktrans) estimates calculated and compared to the ground truth [*K*^trans^](https://osipi.github.io/OSIPI_CAPLEX/quantities/#Ktrans).

### 2.3 Clinical Study

Imaging data acquired for another study^4^ were analysed: 12 ischaemic stroke and 2 transient ischaemic attack survivors scanned chronically (6 – 24 months) after symptom onset, and 20 age-matched controls without history of clinical cerebrovascular disease.

#### 2.3.1 Participants

The stroke group consisted of 11 male and 3 female participants with a mean ± SD age of 68 ± 6 years. The control group consisted of 7 male and 13 female participants with a mean ± SD age of 68 ± 8 years. The clinical study was conducted in full conformity with the current revision of the Declaration of Helsinki, with ethical approval obtained from the National Health Service (North West – Preston Research Ethics Committee).

#### 2.3.2 MRI Acquisition

MRI data were acquired on a Philips 3T Achieva scanner (Manchester Clinical Research Facility) with a 32-channel head coil. A dynamic series of 3D *T*_1_-weighted Fast Field Echo (*T*_1_-FFE; spoiled gradient recalled echo) images were acquired with a [repetition time](https://osipi.github.io/OSIPI_CAPLEX/quantities/#TR) of 2.4 ms, [echo time](https://osipi.github.io/OSIPI_CAPLEX/quantities/#TE) of 0.8 ms, [prescribed excitatory flip angle](https://osipi.github.io/OSIPI_CAPLEX/quantities/#Flip%20angle) of 10 degrees, an in-plane resolution of 1.5 mm x 1.5 mm, 32 contiguous axial slices of 4 mm thickness, and an acquisition time of 7.6 s for each dynamic. 160 dynamic images were acquired over 20 minutes. On the [8^th^](https://osipi.github.io/OSIPI_CAPLEX/quantities/#index) dynamic, a gadolinium-based contrast agent (Gd-DOTA; Dotarem, Guebert, France) bolus was administered using a power injector at a dose of 0.1 mmol/kg and an injection rate of 3 mL/s.

Prior to the dynamic scan, a series of additional 3D *T*_1_-FFE images were acquired at 4 different [excitatory flip angles](https://osipi.github.io/OSIPI_CAPLEX/quantities/#Flip%20angle) (2, 5, 10, and 15 degrees) to calculate the [pre-contrast (native) longitudinal relaxation rate](https://osipi.github.io/OSIPI_CAPLEX/quantities/#R10), [*R*_10_](https://osipi.github.io/OSIPI_CAPLEX/quantities/#R10). Acquisition parameters remained consistent with the dynamic scan except that only eight signal averages were acquired, giving a total acquisition time of 60 s per [flip angle](https://osipi.github.io/OSIPI_CAPLEX/quantities/#Flip%20angle). To correct [*R*_10_](https://osipi.github.io/OSIPI_CAPLEX/quantities/#R10) for *B*_1_ field inhomogeneities and flip angle errors, a pair of actual flip angle mapping^21^ images were also acquired with the same voxel size and coverage as the variable flip angle images. This consisted of a pair of 3D *T*_1_-FFE images with [repetition times](https://osipi.github.io/OSIPI_CAPLEX/quantities/#TR) of 25 ms and 125 ms, respectively, echo time of 5 ms, a [flip angle](https://osipi.github.io/OSIPI_CAPLEX/quantities/#Flip%20angle), *α*, of 60 degrees, and an acquisition time of 117 s.

Structural imaging consisted of a 3D *T*_1_-weighted image with *TR* = 8.4 ms, *TE* = 3.9 ms, flip angle 8 degrees, a resolution of 0.94 mm x 0.94 mm x 1 mm, and an acquisition time of 311 s. For the assessment of white matter lesions, a *T*_2_-weighted FLAIR image was acquired with *TR* = 10 s, an inversion time of 2.75 s, *TE* = 140 ms, an in-plane resolution of 0.69 mm x 0.69 mm with 100 contiguous axial slices of 1.3 mm thickness, and an acquisition time of 450 s.

#### 2.3.3 DCE-MRI Image Processing & Tracer Kinetic Modelling

A *B*_1_-corrected [pre-contrast (native) *R*_1_ map (*R*_10_)](https://osipi.github.io/OSIPI_CAPLEX/quantities/#R10) map was estimated using the [variable flip angle method](https://osipi.github.io/OSIPI_CAPLEX/perfusionProcesses/#VFA) by fitting the [spoiled gradient recalled echo model](https://osipi.github.io/OSIPI_CAPLEX/perfusionModels/#SPGR%20model) described by equation 2 to the signal from each voxel of the variable flip angle images using [non-linear least-squares minimisation](https://osipi.github.io/OSIPI_CAPLEX/generalPurposeProcesses/#NLLS) in Madym^17^ v4.23.0.

The dynamic series of post-contrast *T*_1_-weighted images were aligned using the co-registration function in SPM12 with the first dynamic image as a reference. Using an estimated [*S*_0_](https://osipi.github.io/OSIPI_CAPLEX/quantities/#S_0) from the 6 pre-bolus dynamic images, dynamic estimates of [*R*_1_(*t*](https://osipi.github.io/OSIPI_CAPLEX/quantities/#R1)) were computed for each voxel of the registered dynamic series of *T*_1_-weighted images. The [signal time-course, *S*_t_(*t*)](https://osipi.github.io/OSIPI_CAPLEX/quantities/#S), was thus converted to an [indicator concentration time-course, *C*_t_(*t*)](https://osipi.github.io/OSIPI_CAPLEX/quantities/#C), as described by equation 3, using the [*R*_10_ map](https://osipi.github.io/OSIPI_CAPLEX/quantities/#R10) and the [longitudinal relaxivity](https://osipi.github.io/OSIPI_CAPLEX/quantities/#r1) of the contrast agent, which was assumed to be 3.4 s^-1^mM^-122^. *C*(*t*) measurements were also corrected for errors in the applied flip angle using the flip angle map.

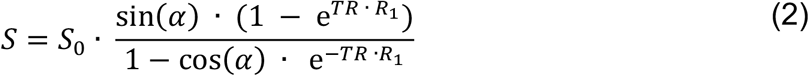

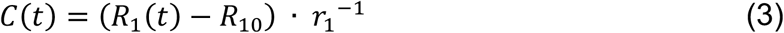

The [indicator concentration in plasma, *C*_p_](https://osipi.github.io/OSIPI_CAPLEX/quantities/#C), was calculated using equation 4:

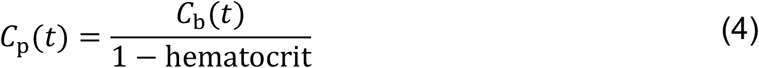

where the [indicator concentration in blood, *C*_b_(*t*)](https://osipi.github.io/OSIPI_CAPLEX/quantities/#C), the vascular input function, was generated using a region of approximately 50 voxels [drawn manually](https://osipi.github.io/OSIPI_CAPLEX/generalPurposeProcesses/#Freehand) within the superior sagittal sinus on the [last dynamic image](https://osipi.github.io/OSIPI_CAPLEX/quantities/#index) in the DCE-MRI series. The superior sagittal sinus has been recommended by consensus guidelines as it is a large vessel with clear post-contrast enhancement, leading to ease of delineation without partial volume effects, and without inflow effects,^23^ and has been shown to reduce measurement variability compared to an arterial input function.^24^ The [mean signal](https://osipi.github.io/OSIPI_CAPLEX/quantities/#S) within this region was extracted for each dynamic to generate [*C*_b_(*t*)](https://osipi.github.io/OSIPI_CAPLEX/quantities/#C). [Hematocrit](https://osipi.github.io/OSIPI_CAPLEX/quantities/#Hct) was assumed to be 0.40 for female participants and 0.45 for male participants.

An Extended Tofts model of indicator exchange (Figure 1a, equation 5) was fit to the [indicator concentration time-course *C*(*t*)](https://osipi.github.io/OSIPI_CAPLEX/quantities/#C) on a voxel-wise basis using constrained [non-linear least-squares minimisation](https://osipi.github.io/OSIPI_CAPLEX/generalPurposeProcesses/#NLLS) in Madym^17^ for 4 parameters: the contrast agent [volume transfer constant across the BBB, *K*^trans^](https://osipi.github.io/OSIPI_CAPLEX/quantities/#Ktrans), the [fractional blood plasma volume, *v*_p_](https://osipi.github.io/OSIPI_CAPLEX/quantities/#v), the [fractional extravascular extracellular volume, v_e_](https://osipi.github.io/OSIPI_CAPLEX/quantities/#v), and *T*_0_. *T*_0_ is the offset time between [*C*_t_(*t*)](https://osipi.github.io/OSIPI_CAPLEX/quantities/#C) and [*C*_p_(*t*)](https://osipi.github.io/OSIPI_CAPLEX/quantities/#C), where the input function is shifted to [*C*_p_(*t* – *T*_0_)](https://osipi.github.io/OSIPI_CAPLEX/quantities/#C) when calculating the predicted tissue concentration [*C*_t_(*t*)](https://osipi.github.io/OSIPI_CAPLEX/quantities/#C) to account for the time delay between the vascular input function and tissue enhancement. A [Patlak model](https://osipi.github.io/OSIPI_CAPLEX/perfusionModels/#Patlak) of contrast agent uptake (Figure 1b, equation 6) was fit for 3 parameters: [*K*^trans^, *v*_p_](https://osipi.github.io/OSIPI_CAPLEX/quantities/#Ktrans), and *T*_0_. An intravascular model was fit similarly to the Patlak model but with [*K*^trans^](https://osipi.github.io/OSIPI_CAPLEX/quantities/#Ktrans) fixed to zero, fitting for only [*v*_p_](https://osipi.github.io/OSIPI_CAPLEX/quantities/#v) and *T*_0_ (Figure 1c, equation 7), essentially scaling and shifting the vascular input function.

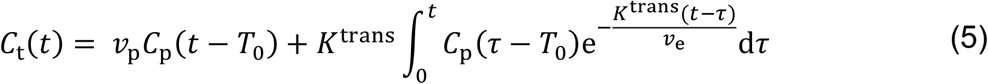

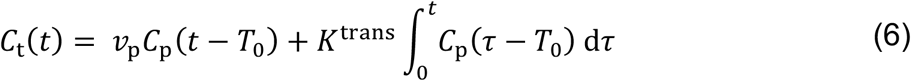

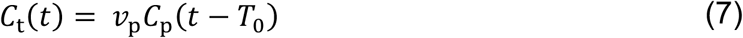

Constraints on the fitted parameters were as follows: between [-1 min^-1^](https://osipi.github.io/OSIPI_CAPLEX/quantities/#MP_L) and [1 min^-1^](https://osipi.github.io/OSIPI_CAPLEX/quantities/#MP_U) for [*K*^trans^](https://osipi.github.io/OSIPI_CAPLEX/quantities/#Ktrans), between 0 and [(1 – hematocrit)](https://osipi.github.io/OSIPI_CAPLEX/quantities/#MP_U) for [*v*_p_](https://osipi.github.io/OSIPI_CAPLEX/quantities/#v), ≤1 for ([*v*_p_](https://osipi.github.io/OSIPI_CAPLEX/quantities/#v) + [*v*_e_](https://osipi.github.io/OSIPI_CAPLEX/quantities/#v)), and between -20 and [20 s](https://osipi.github.io/OSIPI_CAPLEX/quantities/#MP_U) for *T*_0_. Though unphysiological, the negative lower bound for [*K*^trans^](https://osipi.github.io/OSIPI_CAPLEX/quantities/#Ktrans) is necessary to avoid positive bias in noisy [*K*^trans^](https://osipi.github.io/OSIPI_CAPLEX/quantities/#Ktrans) values, which may be very close to zero in healthy tissue. The negative lower bound on *T*_0_ allows for the possibility that the regional blood circulation may occur later than the sagittal sinus peak concentration. The residuals of the first 15 time-points were ignored in the voxel-wise model fitting, with each model fit to the remaining 145 time-points. Optimisation of this cutoff time is presented in the Supplementary Materials.

#### 2.3.4 Model Selection

The [Akaike information criterion](https://osipi.github.io/OSIPI_CAPLEX/quantities/#AIC)^25^ [*(AIC)*](https://osipi.github.io/OSIPI_CAPLEX/quantities/#AIC) was used to determine the best-fitting model. The *AIC* was computed for each model, m, using the residual sum of squared errors on the fit, *RSS*, and the number of free parameters, *K*, as set out in equation 8, where *N* corresponds to the number of data points (dynamic images). The Akaike Weight (*AW*) was calculated from the *AIC*s as described by equation 9 and the model with the largest *AW* selected as the best-fitting model for each voxel. [*K*^trans^](https://osipi.github.io/OSIPI_CAPLEX/quantities/#Ktrans) estimates from the best-fitting model were selected for each voxel to produce best-fitting ‘model selection’ [*K*^trans^](https://osipi.github.io/OSIPI_CAPLEX/quantities/#Ktrans) maps.

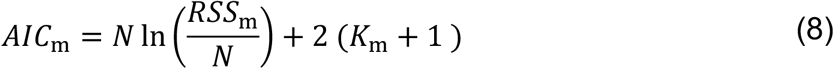

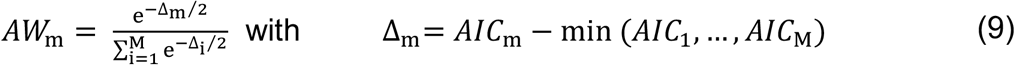

In voxels where the estimated [*v*_p_](https://osipi.github.io/OSIPI_CAPLEX/quantities/#v) < 0.1%, the tissue was presumed to be poorly perfused and no model or [*K*^trans^](https://osipi.github.io/OSIPI_CAPLEX/quantities/#Ktrans) estimate was considered for these voxels.

#### 2.3.5 Post-Processing & Region of Interest Analysis

White matter hyperintensities were defined from the *T*_2_-weighted FLAIR images using the lesion segmentation toolbox (LST)^26^ in SPM12 with the 3D *T*_1_-weighted image as a reference for co-registration. Regions of interest corresponding to the cerebral cortex, cerebral NAWM, and DGM were segmented on the 3D *T*_1_-weighted image using FreeSurfer v7.3.2’s ‘recon-all’ and voxels overlapping with the WMH region removed. The 3D *T*_1_-weighted image was co-registered to the [first dynamic image](https://osipi.github.io/OSIPI_CAPLEX/quantities/#index) of the DCE-MRI acquisition using the co-registration function in SPM12 and the transformation applied to all regions of interest. All co-registered regions were visually checked for alignment with the DCE-MRI dynamic series.

The second aim of this study was to characterise the spatial distribution of best-fitting models across the cortex, DGM, NAWM, and WMH in stroke survivors and controls. To address aim 2, the number of voxels selecting each model was determined as a percentage of the total number of voxels for each region. The [median value](https://osipi.github.io/OSIPI_CAPLEX/generalPurposeProcesses/#CalcMedian) of [*K*^trans^](https://osipi.github.io/OSIPI_CAPLEX/quantities/#Ktrans) was computed for each region and each model, as well as for the model selection [*K*^trans^](https://osipi.github.io/OSIPI_CAPLEX/quantities/#Ktrans) map (i.e., a map of the [*K*^trans^](https://osipi.github.io/OSIPI_CAPLEX/quantities/#Ktrans) from the best-fitting model at each voxel).

For illustrative purposes, small regions of approximately 100 voxels were hand drawn within the cortex, NAWM, DGM and WMH regions of a representative control and stroke brain. The mean estimated concentration from the intravascular, [Patlak model](https://osipi.github.io/OSIPI_CAPLEX/perfusionModels/#Patlak) and [Extended Tofts model](https://osipi.github.io/OSIPI_CAPLEX/perfusionModels/#ETM) were extracted and the residuals plotted in order to demonstrate examples of the model performance. To demonstrate patterns across the groups, all maps were co-registered to the *T*_1_-weighted image and normalised to MNI space using the ‘normalise’ function in SPM12, and group-average model and [*K*^trans^](https://osipi.github.io/OSIPI_CAPLEX/quantities/#Ktrans) maps produced for both the control and stroke groups.

#### 2.3.6 Statistics

Mann Whitney tests were used to test for differences in the proportion of voxels selecting the intravascular model as best-fitting between the control and stroke group for the cortex, DGM and NAWM. This was repeated for the Patlak and Extended Tofts model.

The third aim of this study was to compare *K*^trans^ estimates derived from voxel-wise model selection to those from conventional Patlak fitting. To address aim 3, we tested for differences between [median values](https://osipi.github.io/OSIPI_CAPLEX/generalPurposeProcesses/#CalcMedian) of [Patlak model](https://osipi.github.io/OSIPI_CAPLEX/perfusionModels/#Patlak) and model selection [*K*^trans^](https://osipi.github.io/OSIPI_CAPLEX/quantities/#Ktrans) estimates in the four regions of interest using Wilcoxon signed-rank tests.

Finally, aim 4 was to assess whether voxel-wise model selection better distinguishes pathological from normal-appearing tissue compared to conventional Patlak fitting. To address aim 4, Wilcoxon signed-rank (paired) tests were used to test for differences between [median values](https://osipi.github.io/OSIPI_CAPLEX/generalPurposeProcesses/#CalcMedian) of [*K*^trans^](https://osipi.github.io/OSIPI_CAPLEX/quantities/#Ktrans) in the NAWM and WMH of stroke patients for the [Patlak model](https://osipi.github.io/OSIPI_CAPLEX/perfusionModels/#Patlak) and model selection method. Mann-Whitney (unpaired) tests were used to test for differences between [median values](https://osipi.github.io/OSIPI_CAPLEX/generalPurposeProcesses/#CalcMedian) of [*K*^trans^](https://osipi.github.io/OSIPI_CAPLEX/quantities/#Ktrans) in controls and stroke patients for the [Patlak model](https://osipi.github.io/OSIPI_CAPLEX/perfusionModels/#Patlak) and model selection method.

## 3 Results

### 3.1 Simulations

A heatmap of the ground truth *K*^trans^ for the array of simulated concentration time curves is presented in Figure 2a. The best-fitting model was largely dependent on the magnitude of *K*^trans^, with the Extended Tofts model chosen for the high permeability regime (>10^−3^ min^-1^), the Patlak model chosen in the subtle permeability regime (10^−4^ – 10^−3^ min^-1^), and the intravascular model selected at 10^−4^ min^-1^, as shown in Figure 2b. At low CNR, the intravascular model was best-fitting for voxels with up to *K*^trans^ ∼ 3 × 10^−4^ min^-1^, and Patlak up to *K*^trans^ ∼ 6 × 10^−3^ min^-1^.

**Figure 2.**
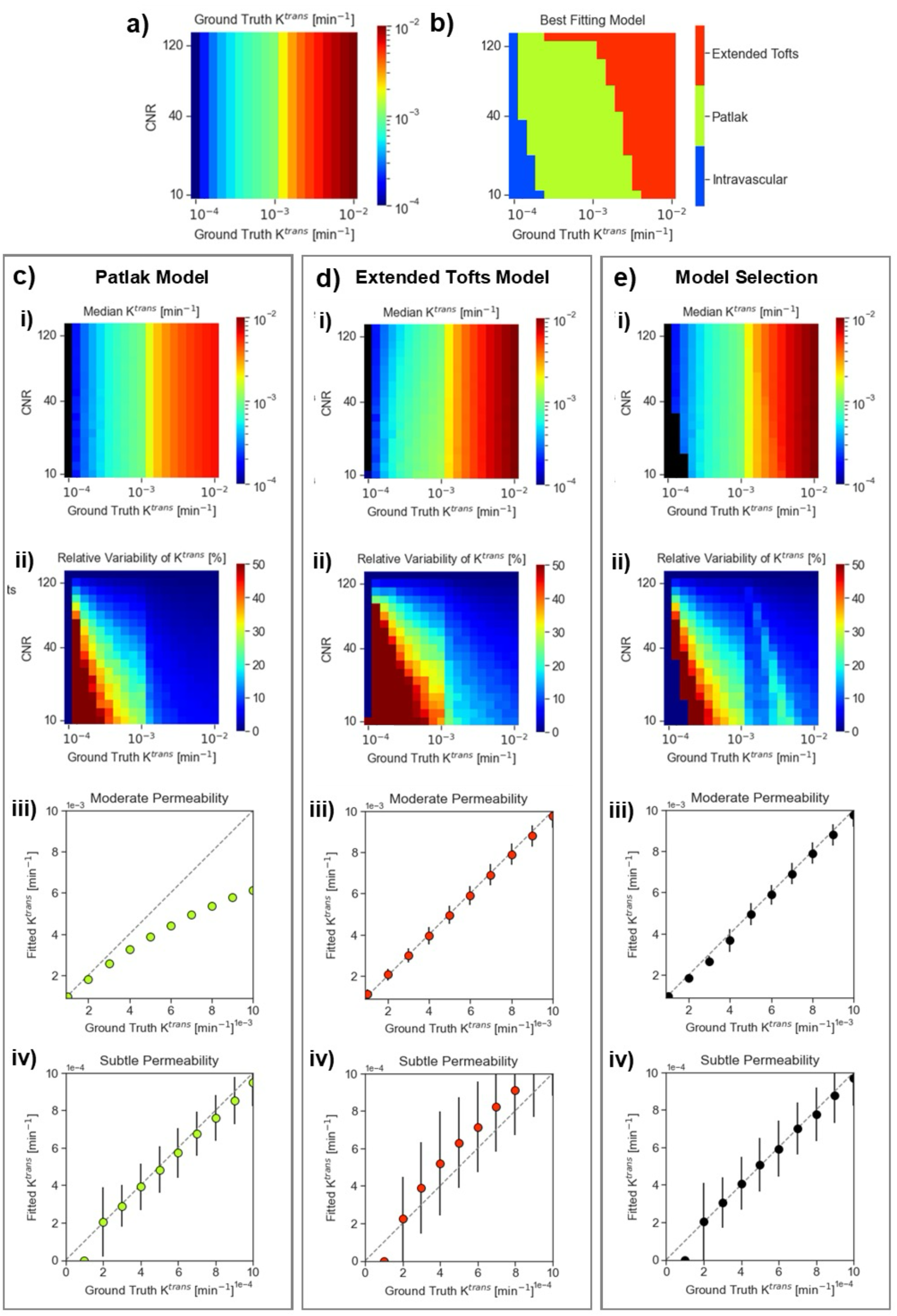
Model Selection Output for Simulated ‘White Matter’. **(a)** Heatmap of the ground truth *K*^trans^ input for the simulated concentration time curves. **(b)** Best-fitting model map for varying ground truth *K*^trans^ and CNR. **(c – e)** For **(c)** the Patlak model, **(d)** the Extended Tofts Model, **(e)** Model selection: **(i)** Heatmap of median fitted *K*^trans^ estimates at varying ground truth *K*^trans^ and CNR. **(ii)** Heatmap of relative variability (median absolute deviation as a percentage of the median) of *K*^trans^ estimates at varying ground truth *K*^trans^ and CNR. **(iii - iv)** Median fitted *K*^trans^ is plotted against ground truth or a CNR roughly corresponding to that of control white matter voxels (CNR ∼ 40) for the (iii) high permeability and (iv) subtle permeability regimes. Error bars shown are the median absolute deviation.

At a CNR equivalent to that of white matter voxels, median Patlak measures of *K*^trans^ substantially underestimated *K*^trans^ in the high permeability regime but provide good estimates for subtle permeability (Figure 2c). Median Extended Tofts measures of *K*^trans^ provided a good estimate of *K*^trans^ at both high and subtle permeability, but with more variability and hence poorer precision than Patlak in the subtle leakage regime (Figure 2d). The median model selection method measures of *K*^trans^ accurately estimated *K*^trans^ across the subtle and high permeability regimes (Figure 2e) with lower variability and hence better precision than the Extended Tofts model at small *K*^trans^. The Patlak, Extended Tofts, and model selection methods all estimated a median *K*^trans^ of zero at a ground truth *K*^trans^ ∼ 10^−4^ min^-1^ indicating that this is the lower limit of detectability at typical voxel-level CNR. Experiments independently varying *K*^trans^ with *F*_p_, *v*_p_, *v*_e_, and *T*_10_, as well as Monte Carlo simulations, yielded similar findings and are described in detail in the Supplementary Materials.

### 3.2 Clinical Study

#### 3.2.1 Qualitative Assessment of Voxel-Wise Model Fits

An example ‘best-fitting’ model selection map, model selection *K*^trans^ map, and model fits for 100 voxel tissue samples of a representative stroke participant are presented in Figure 3. The model fits were comparable in the DGM (Figure 3c), estimating a median Patlak model *K*^trans^ = 2.40 × 10^−5^ min^-1^, Extended Tofts model *K*^trans^ = 4.71 × 10^−4^ min^-1^, and model selection *K*^trans^ = 0 min^-1^. In the NAWM, the Patlak model performed adequately, with estimated median Patlak model *K*^trans^ = 1.14 × 10^−4^ min^-1^, Extended Tofts model *K*^trans^ = 9.08 × 10^−4^ min^-1^, and model selection *K*^trans^ = 2.03 × 10^−4^ min^-1^ (Figure 3d).

**Figure 3.**
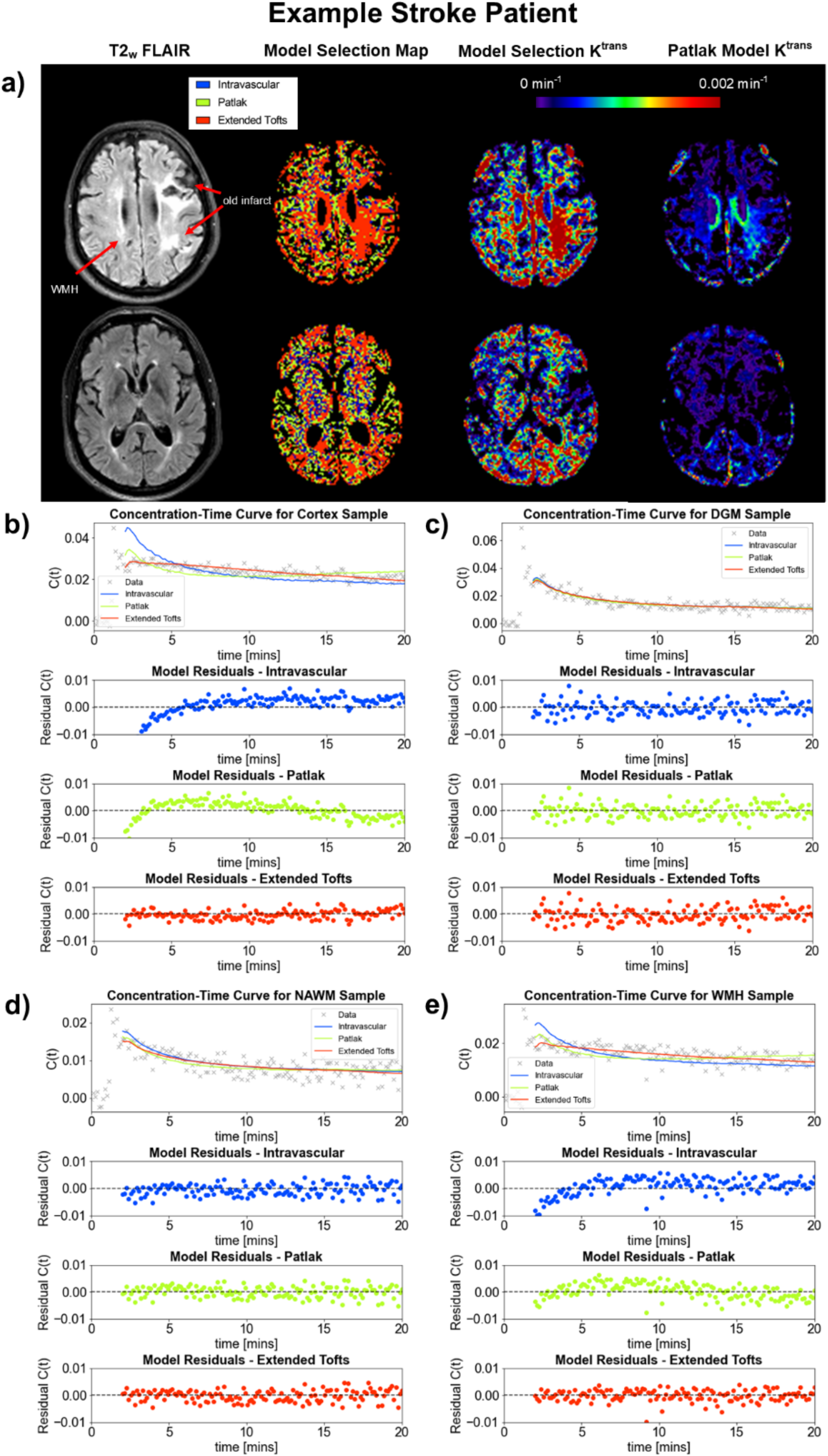
Model Selection Output for an Example Stroke Patient. **(a)** Representative images for a 54-year-old male scanned 1 year after onset of ischaemic stroke. This individual presented with expressive dysphasia and right facial paresis and was diagnosed with an acute left middle cerebral artery territory infarct. A structural *T*_2_-weighted FLAIR image, a map of the selected model for each voxel, best-fitting *K*^trans^ map, and Patlak *K*^trans^ map are shown. The old infarct is indicated on the FLAIR image with a red arrow. **(b)** An example concentration-time curve for approximately 100 voxels in the cortex, median model selection *K*^trans^ = 3.01 × 10^−3^ min^-1^. **(c)** An example concentration-time curve for approximately 100 voxels in the deep grey matter, median model selection *K*^trans^ = 0 min^-1^. **(d)** An example concentration-time curve for approximately 100 voxels in the normal appearing white matter, median model selection *K*^trans^ = 2.03 × 10^−4^ min^-1^. **(e)** An example concentration-time curve for approximately 100 voxels in a white matter hyperintensity region, median model selection *K*^trans^ = 1.66 × 10^−3^ min^-1^. For each graph b – e, the average signal-derived concentration time course is plotted with the fitted concentration time course for the three models, with the respective residuals on the fit for each model shown below.

The intravascular model and Patlak model performed poorly compared to the Extended Tofts model in the cortex (Figure 3b), estimating a median Patlak model *K*^trans^ = 8.14 × 10^−4^ min^-1^, Extended Tofts model *K*^trans^ = 3.05 × 10^−3^ min^-1^, and model selection *K*^trans^ = 3.01 × 10^−3^ min^-1^. Finally, in the WMH sample, the intravascular and Patlak models performed poorly compared to the Extended Tofts model, with estimated median Patlak model *K*^trans^ = 3.46 × 10^−4^ min^-1^, Extended Tofts model *K*^trans^ = 1.82 × 10^−3^ min^-1^, and model selection *K*^trans^ = 1.66 × 10^−3^ min^-1^ (Figure 3e).

#### 3.2.2 Regional Distribution of Best-fitting Models

All models were selected for some proportion of voxels in all regions, however for the stroke group the Extended Tofts model was generally selected more often than for the control group, demonstrated in the group-averaged model selection maps in Figure 4a-b. In the cortex, the Extended Tofts model was selected in a significantly greater proportion of voxels in the stroke group compared to controls (39% vs. 28%, P < 0.0003), Figure 4c-d. The Patlak model was best fitting in significantly fewer voxels (30% vs. 38%, P = 0.015), as was the intravascular model (12% vs. 16%, P < 0.002). Similarly, in the DGM and NAWM the Extended Tofts model was selected in a significantly larger proportion of voxels in the stroke group compared to controls (21% vs. 8%, P < 0.0001 for DGM; 28% vs. 18%, P < 0.0001 for NAWM), while there were not differences in the proportion of voxels preferring the Patlak model (41% vs. 37%, P = 0.07 for DGM; 39% vs. 32%, P = 0.09 for NAWM). The intravascular model was best fitting in significantly fewer voxels in the stroke group compared to controls in the DGM and NAWM (31% vs 38%, P < 0.004 for DGM; 20% vs 27%, P < 0.002 for NAWM). In the WMH, a median of 15% of voxels selected the intravascular model as best-fitting, 25% selected the Patlak model, and 44% of voxels selected the Extended Tofts model.

**Figure 4.**
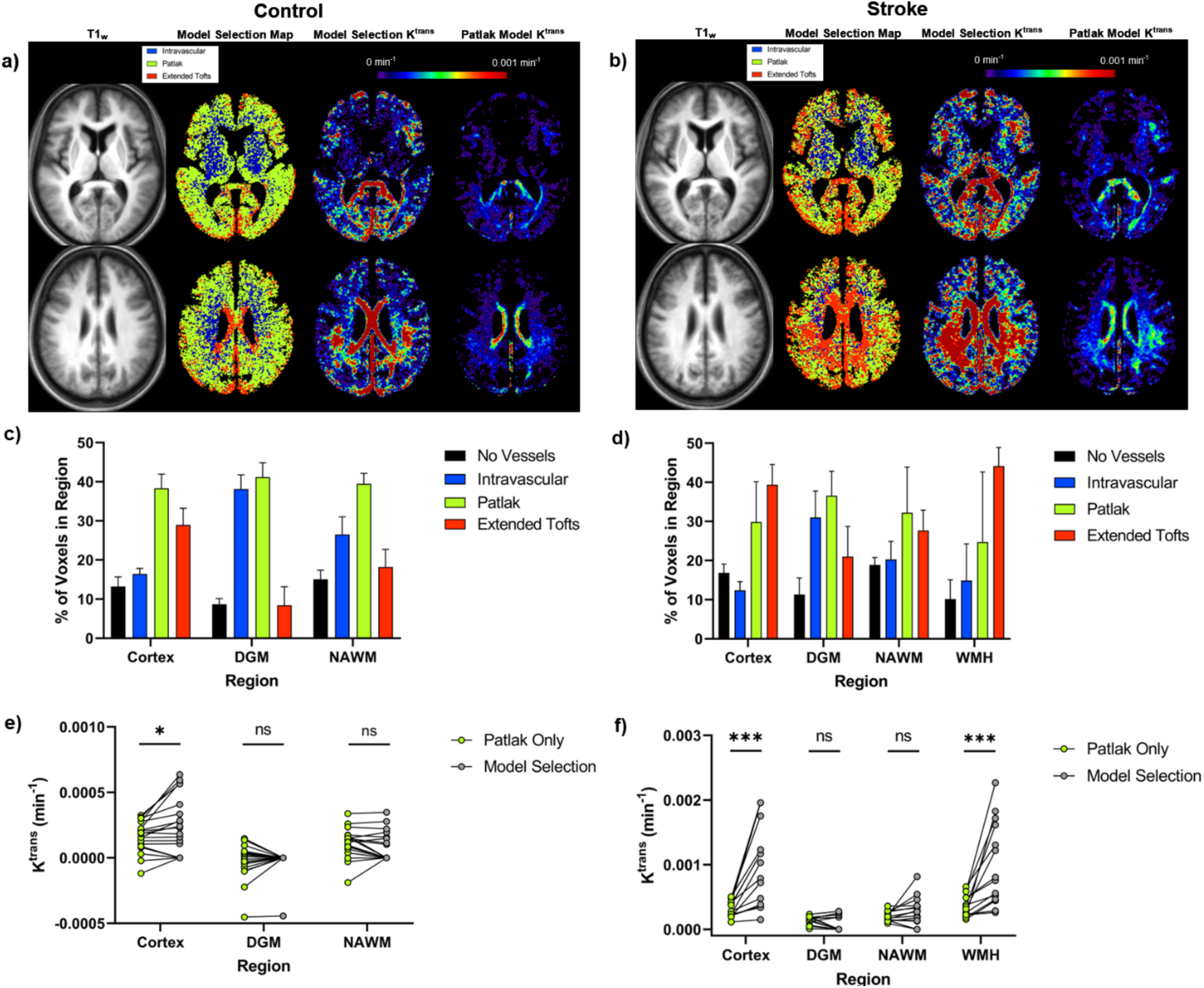
Model Selection Patterns Across Groups. **(a – b)** Group-averaged images for the control (a) and stroke (b) group. A structural *T*_1_-weighted image, a map of the selected model for each voxel, best-fitting *K*^trans^ map, and Patlak *K*^trans^ map are shown. **(c – d)** Median proportion of voxels selecting each model in (c) the cortex, deep grey matter, and normal appearing white matter of controls, and (d) the cortex, deep grey matter, normal appearing white matter, and white matter hyperintensities of stroke patients. Error bars shown are the 95% confidence interval. **(e – f)** Median *K*^trans^ values extracted from the ‘best-fitting’ model selection *K*^trans^ map are compared to median *K*^trans^ values from the conventional Patlak model for (e) the cortex, deep grey matter, and normal appearing white matter of controls, and (f) the cortex, deep grey matter, normal appearing white matter, and white matter hyperintensities of stroke patients. Statistical significance indicated on the plots correspond to results of the Wilcoxon signed-rank tests, where ns corresponds to P ≥ 0.05, * P < 0.05, *** P < 0.001.

#### 3.2.3 Impact on Parameter Estimates

In the control group, median model selection *K*^trans^ estimates were significantly greater than Patlak model *K*^trans^ estimates in the cortex (P = 0.03) but not the DGM (P = 0.9) or NAWM (P = 0.05), shown in Figure 4e. In the stroke group, median model selection *K*^trans^ estimates were significantly greater than Patlak model *K*^trans^ estimates in the cortex (P = 0.0001) and WMH (P = 0.0001), but not the DGM (P = 0.2) or NAWM (P = 0.09). In the stroke group, median model selection *K*^trans^ estimates had greater inter-participant variability in the cortex and WMH compared to estimates from the Patlak model only (Figure 4f). In the control DGM, the model selection *K*^trans^ had markedly less inter-participant variability, with median values converging at zero.

#### 3.2.4 Impact on Interpretation of Parameter Estimates

The absolute differences between *K*^trans^ estimates in the control and stroke groups were greater with the model selection method than the Patlak model only, especially in the cortex. However, the outcome (significant or not) of statistical comparisons was the same for Patlak or model selection (Figure 5a-b). Similarly, differences between *K*^trans^ estimates in the NAWM and WMH of stroke patients were greater with the model selection method than Patlak only, but the outcome of statistical comparisons were the same (Figure 5c-d).

**Figure 5.**
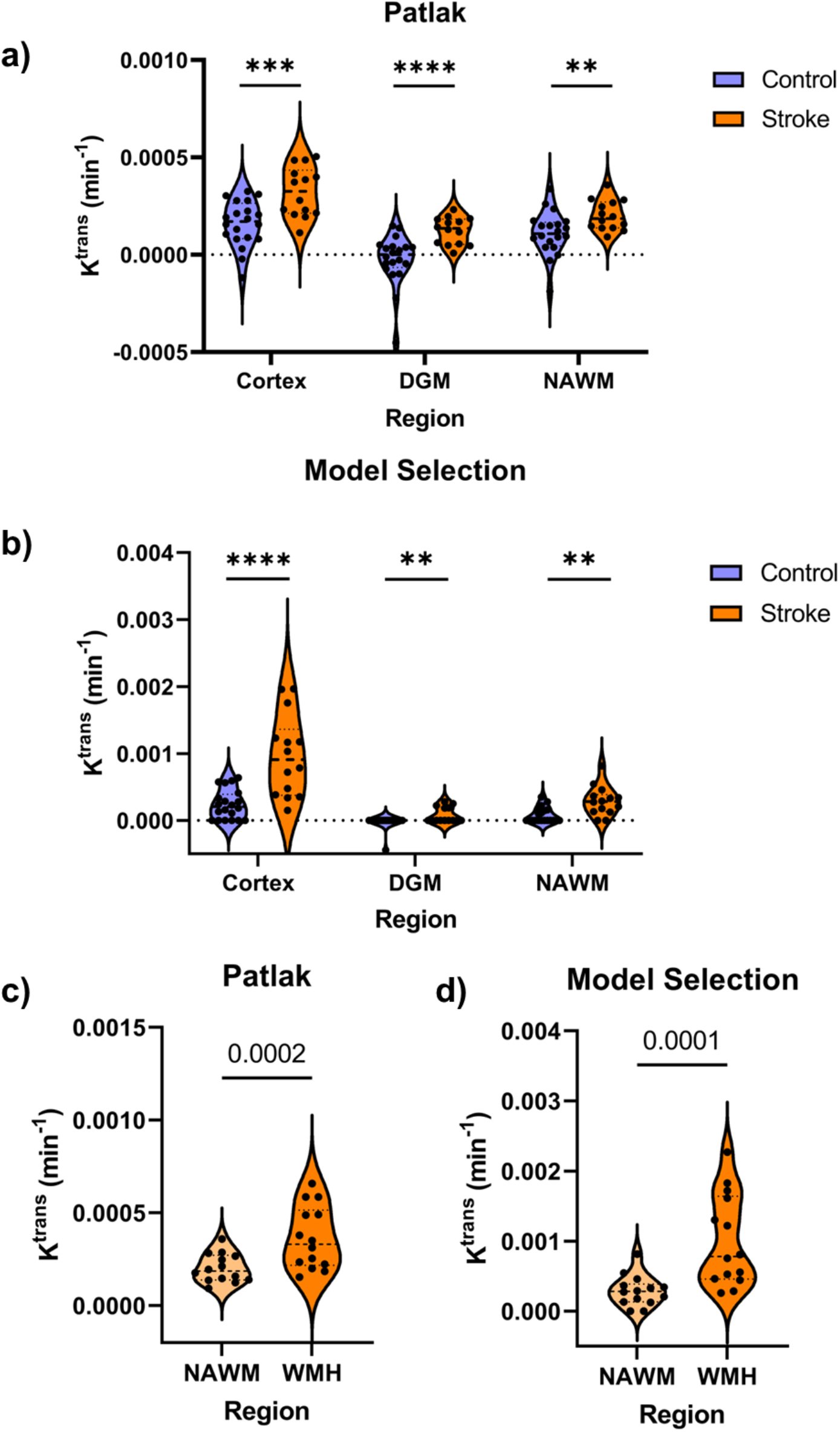
Impact on Interpretation of Results. **(a-b)** Comparison of median *K*^trans^ values between the control and stroke group for the cortex, deep grey matter, and normal appearing white matter regions for the Patlak model (a) and the model selection method (b). **(c-d)** Comparison of median *K*^trans^ values between normal appearing white matter and white matter hyperintensities of stroke patients for the Patlak model (c) and the model selection method (d). Statistical significance indicated on the plots correspond to results of the statistical comparisons (Mann-Whitney tests for a – b and Wilcoxon signed-rank tests for c – d), where ** corresponds to P < 0.01, *** P < 0.001, **** P < 0.0001. *Please note differences in axes scale for each plot*.

## 4 Discussion

This study assessed the utility of voxel-wise tracer kinetic model selection for the assessment of BBB breakdown in neurological disease using simulated and *in-vivo* DCE-MRI data from chronic stroke survivors and controls. We demonstrate that the Patlak model underestimates *K*^trans^ in high permeability voxels (≥ 10^−3^ min^-1^), and that model selection produces more accurate quantification of a wider range of *K*^trans^ values than any single model used alone. All three models were selected in a proportion of brain voxels, with the Extended Tofts model preferred in a large proportion of cortical voxels in both stroke survivors and controls, as well as within the WMH, resulting in significantly greater *K*^trans^ estimates compared to Patlak fitting alone. Though differences in *K*^trans^ between patients and controls, and pathological versus ‘normal appearing’ tissue, were larger with the model selection method than Patlak fitting alone, the interpretation of statistical tests of differences were similar for both approaches.

Simulations confirmed that the Extended Tofts model provides substantially greater accuracy compared to the Patlak model for *K*^trans^ estimation at high permeabilities. This is because the Extended Tofts model allows for back-flux of the contrast agent from the extravascular space, which becomes necessary when BBB leakage is severe enough for contrast agent clearance to occur within the scan duration, invalidating the Patlak model’s assumption of slow, linear leakage of contrast agent. The use of a sub-optimal model in such voxels risks systematic underestimation of the true BBB permeability. Simulations additionally showed that the simpler Patlak model should be adequate in the case of more subtle permeabilities and was associated with a lower variability in noisy estimates compared to the Extended Tofts model. Similar results have been shown in a study comparing the Patlak and Extended Tofts model in multiple sclerosis,^27^ supporting the generalisability of these findings beyond stroke.

The intravascular model serves an important role in the model selection framework, capturing voxels where no detectable enhancement above noise is present. Inclusion of the intravascular model reduced the incidence of physiologically implausible negative *K*^trans^ estimates, a known artefact of fitting models to the high levels of noise in voxel DCE-MRI data,^23,28^ as evident in Figures 4 and 5. Simulations showed that the intravascular model was most frequently selected at *K*^trans^ ∼ 10^−4^ min^-1^, suggesting this is the lower limit of detectability in our noisy voxel-wise data. Though the model selection performance was relatively robust to noise, at very high noise levels there was increasing bias toward the intravascular model, and factors such as patient motion, low SNR, and scanner drift may similarly obscure subtle signal enhancement, leading to an underestimation of the true *K*^trans^. It is therefore important to note that if the intravascular model is selected, this may be a result of technical limitations rather than the BBB being truly closed. High noise may also increase variance in the Akaike Weights themselves, potentially reducing precision compared to a single model approach.^29^ Detection of very subtle BBB leakage may be more achievable with regional analysis where the CNR is much greater than that of voxel-wise data.

In the clinical data, the spatial distribution of preferred models was heterogenous, even within controls. The conventionally used Patlak model was largely the preferred model in the DGM, NAWM, and cortex voxels of controls, though the Extended Tofts model was chosen in 30% of voxels in the cortex. The intravascular model was not the dominantly preferred model in any region, though best described 38% of the deep grey matter voxels of controls. Though it is known that the BBB is not homogenous across the brain,^14^ and that BBB disruption occurs in normal aging,^30,31^ there is little evidence with regard to BBB differences between DGM and the cortex in the healthy older human brain. Here we have a relatively large voxel size of 1.5 × 1.5 × 4 mm, and it is therefore possible that there are partial volume effects with some cortical voxels and the meninges, vessels within which are more permeable than those in the parenchyma,^32^ and this could partially explain our finding.

In stroke survivors, the Patlak model was the preferred model in the DGM and NAWM. In the cortex and WMH, the Extended Tofts model was preferred in the largest proportion of voxels. This resulted in significantly greater average *K*^trans^ estimates, larger differences between stroke survivors and controls, as well as between NAWM vs. WMH, though the interpretation of statistical tests of differences were the same with Patlak or the model selection method. Elevated *K*^trans^ in the WMH compared to NAWM 3+ months after mild stroke has been previously reported using DCE-MRI with Patlak modelling,^33^ as well as differences between *K*^trans^ in stroke/TIA patients and age-matched controls.^34^ Our finding suggests that *K*^trans^ may be underestimated when the conventional Patlak model alone is used in this patient group, and that BBB impairment chronically after ischaemic stroke could be more severe than previously reported. We also find a greater range and variability in model selection *K*^trans^ estimates compared to Patlak estimates, and this effect is more pronounced in the stroke group than the controls, possibly by more accurately reflecting the biological variability in BBB damage caused by stroke in this group.

Previous work by Heye et al.^13^ has shown that the Patlak model fits best in the DGM, NAWM, and WMH of control and mild stroke survivors, however this study performed modelling on a regional basis as opposed to voxel-wise, and with an acquisition protocol of much slower temporal resolution than is used here (73 s vs 7.6 s). While slow temporal resolution is adequate for the detection of subtle tracer leakage,^13^ faster tracer dynamics in tissue with a more permeable BBB may require higher temporal resolution for accurate *K*^trans^ estimation. The previous study also used a lower field strength (1.5T vs 3T), which may affect SNR and therefore the choice of model. Additionally, we were unable to explicitly exclude potential old infarct tissue from our definition of WMHs, whereas the previous study separated the two, which could contribute to some of the differences in findings.

The principal strength of the voxel-wise model selection method in DCE-MRI is the ability to more accurately estimate a greater range of *K*^trans^ values in a single brain, revealing more pronounced differences between normal tissue and pathology, as we have shown in this study. In cases such as brain tumour or stroke, where regions of both extreme and subtle BBB pathology are expected and of interest, this could be particularly useful. Furthermore, model selection maps may provide further information about the underlying BBB physiology than *K*^trans^ maps alone. In cases where the Extended Tofts model fits best, the BBB may be very leaky, whereas regions selecting the intravascular model are likely to be healthy tissue. Thus, knowledge of the voxel-wise distribution of preferred models could be used to infer the volume and location of leaky tissue, which could be especially valuable for determining where in the brain BBB leakage may be occurring in pathologies without a specific lesion of interest, such as dementia. Additionally, where the Extended Tofts model is selected, the extravascular extracellular volume fraction, *v*_e_, is additionally estimated. Importantly, model selection reduces errors in *v*_e_ that would arise from fitting the Extended Tofts model to the entire brain volume, since *v*_e_ can only be estimated reliably in voxels where detectable clearance occurs. Estimates of *v*_e_ may be able to offer valuable additional information about the brain after stroke, where alterations in the relative volume of extravascular extracellular space could indicate oedema or tissue loss.

Quantitative DCE-MRI analysis methods are already difficult to adopt outside a research context and the magnitude of derived parameters such as *K*^trans^ can vary between sites based on the chosen analysis method.^35^ This has been attributed in part to the several decision steps required to form an analysis pipeline, such as which model to use, the optimisation strategy used in fitting, etc.^36^ Though more computational resources may be necessary to perform the model selection method than the fitting of a single model, the decision of which model to use is data-driven and thus may be a step towards a more robust and reproducible pipeline if adopted. Further to this, for the modelling implementation we have used an open-source, publicly available software^17^ with demonstrated reproducibility in the Open Science for Perfusion Imaging DCE-MRI challenge.^35^ Accurate, reproducible quantification of spatially heterogeneous BBB dysfunction is important for the study of neurological disease, and model selection should be adopted as a component of voxel-wise DCE-MRI analysis pipelines to achieve this.

## 5 Data Availability Statement

Data are available upon reasonable request. Imaging processing pipeline can be accessed via GitHub: https://tinyurl.com/3wbswtm4.

## 6 Acknowledgements

MRI scan costs were supported by The Sydney Driscoll Neuroscience Foundation and the United Kingdom Research and Innovation Engineering and Physical Sciences Research Council (grant number EP/M005909/1). OAJ would like to acknowledge support from the United Kingdom Research and Innovation Engineering and Physical Sciences Research Council doctoral training programme at the University of Manchester, the Stroke-IMPaCT network (Leduqc Foundation, grant number 19CVD01), and the StrokeCog-BBB study (National Institute of Neurological Disorders and Stroke, National Institutes of Health, grant number R01NS124927).

