## Supplementary Materials for "Voxel-wise tracer kinetic model selection for DCE-MRI measurements of blood-brain barrier leakage"

#### 1 Determining the optimal cutoff time for model fitting.

The models described by equations 5 – 7 are associated with several assumptions that are violated during the first pass of the contrast agent, leading to errors in parameter estimation. These violated assumptions include: compartments that are not well mixed, effects of flow, and errors in the accuracy of the VIF unless temporal resolution is very high, due to the rapid influx of contrast agent. For the Patlak model especially, inclusion of these early time points has been shown to lead to errors in parameter estimation.[1] However, early time-points after the first pass may be important for the detection of more substantial contrast agent leakage. Therefore, we aimed to exclude the most problematic first-pass effects while retaining sufficient data to accurately characterise a range of tracer kinetic behaviours.

A Patlak plot consists of plotting the ratio of $\mathrm{C_t(t)/C_p(t)}$ against $\int \mathrm{C_p(\tau)\,d\tau/C_p(t)}$, where perfect compliance with the model assumptions would yield a straight line, and deviations from linearity indicate violations of the model assumptions, which are particularly evident in the first pass. We generated Patlak plots from the regional grey/white matter data for each participant to assess the impact of these early time points on errors in the linear assumption of the Patlak model and determine the optimal amount of early time-point data to exclude. Example Patlak plots with and without the early time points removed, as well as plots of the mean squared error (MSE) on the linear fit as a function of cutoff time, are presented in Supplementary Figure 1. Informed by this analysis, we selected a cutoff time after the stabilisation of the MSE metric across participants of around 1 minute after injection, and the residuals of the first 15 time-points were ignored in the voxel-wise model fitting, with each model fit to the remaining 145 time-points.

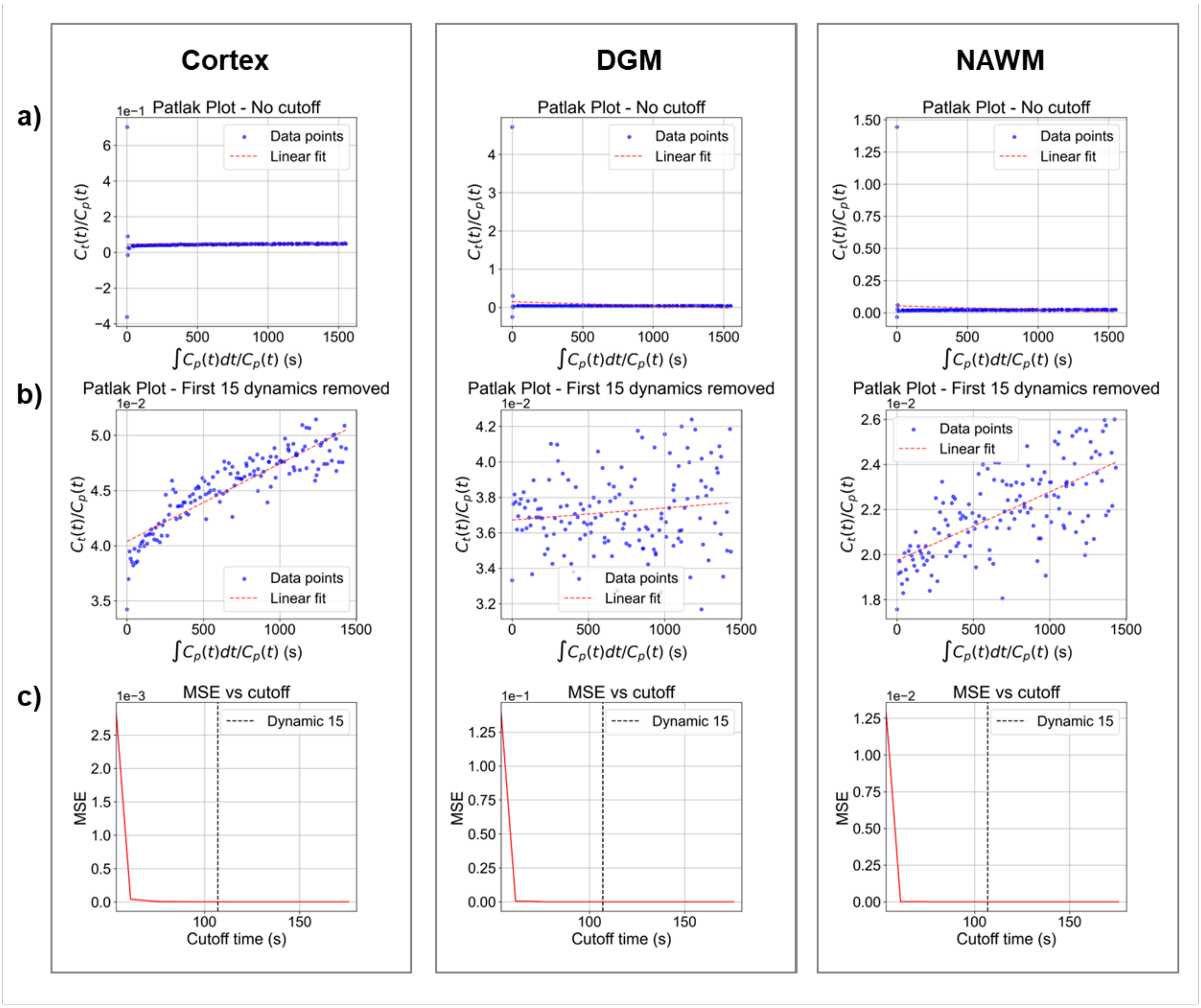


**Supplementary Figure 1. Patlak Plots For Determining Optimal Cutoff Time.** All plots are from an example control participant. Averaged over the cortex, deep grey matter (DGM) and normal-appearing white matter (NAWM) regions of interest, respectively: **(a)** A Patlak plot for the entire time-series. **(b)** A Patlak plot with one minute removed post-contrast. **(c)** Mean squared error (MSE) on the Patlak plot fitting as a function of cutoff time, with a vertical dashed line indicating the early time points excluded from the model fitting.

### 2 Exploration of parameter space using simulated data.

Using the simulation framework described in the main methods, we further explored parameter space to ensure the generalisability of our finding that the selected 'best-fitting' model is determined primarily by the magnitude of the ground truth *PS*, and to ensure robust fitting across the full range of physiologically plausible parameter combinations in both health and disease.

First, we systematically varied the ground truth PS with each individual parameter at a fixed level of noise representative of voxel-level DCE-MRI data in our participants. Parameters were varied across the ranges described in Supplementary Table 1 and the data simulation and model fitting/selection performed as described in the main methods. For each experiment, the non-varying parameters were fixed to the following values: $F_p$ = 0.4 min$^{-1}$, $v_p$ = 2%, $v_e$ = 20%, $T_{10}$ = 1000 ms.

**Supplementary Table 1.** Input values and physiological representations for simulations independently varying individual parameters.

| Input parameter | Units | Input values | Physiological Representation of Range |
|---|---|---|---|
| Permeability surface area product, *PS* | min$^{-1}$ | 0.0001, 0.0005, 0.001, 0.005, 0.01 | Subtle to very leaky BBB. |
| Plasma flow, $F_p$ | min$^{-1}$ | 0.1, 0.3, 0.6, 0.8, 1.2 | Hypoperfused/ischaemic to highly perfused tissue. |
| Plasma volume fraction, $v_p$ | % | 1%, 2%, 4%, 6%, 10% | Weakly vascularised to highly vascularised tissue. |
| Extravascular extracellular volume fraction, $v_e$ | % | 5%, 10%, 15%, 20%, 25% | Small interstitial space (cell swelling/oedema) to large interstitial space (atrophy). |
| Baseline T1, $T_{10}$ | ms | 800, 1000, 1250, 1500, 2000 | Underlying brain tissue type (WM to blood). |

The effect of variable $F_p$ on the accuracy of $K^{trans}$ estimation is shown in Supplementary Figure 2. In general, findings were comparable to the use of a fixed $F_p$ = 0.4 min$^{-1}$ representative of normal white matter, the selection of the preferred model was driven largely by the magnitude of the ground truth *PS*, and the 'best-fitting' $K^{trans}$ estimates had the highest accuracy compared to any of the models used alone. Only the hypoperfused scenario ($F_p$ = 0.1 min$^{-1}$) led to a small negative bias on $K^{trans}$ estimates compared to normal or high perfusion. When plasma flow is high relative to permeability, the contrast agent delivery to the tissue is limited by permeability and $K^{trans} \approx PS$. Similarly, when flow is low relative to permeability, $K^{trans} \approx F_p$. In a mixed regime, the measured $K^{trans}$ is an effective combination of both plasma flow and vessel permeability.[2] Thus, this bias may indicate the limit of the assumption that $PS >> F_p$ has been reached, suggesting the beginning of the mixed regime whereby the measured $K^{trans}$ is a combination of both plasma flow and vessel permeability. Similarly to the fixed-$F_p$ simulation experiment, the model selection $K^{trans}$ estimates had the highest accuracy compared to any of the individual models used alone.

The effect of variable $v_p$ on the accuracy of $K^{trans}$ estimation is shown in Supplementary Figure 3. In general, findings were comparable to the use of a fixed $v_p$ = 2% representative of normal white matter, the selection of the preferred model was driven largely by the magnitude of the ground truth $PS$, and the 'best-fitting' $K^{trans}$ estimates had the highest accuracy compared to any of the models used alone. The highly vascularised scenario ($v_p$ = 10%) led to underestimation of $K^{trans}$ compared to the normal range ($v_p$ = 2 – 6%).

The effect of variable $v_e$ on the accuracy of $K^{trans}$ estimation is shown in Supplementary Figure 4. In general, findings were comparable to the use of a fixed $v_e$ = 20%, representative of normal white matter, the selection of the preferred model was driven largely by the magnitude of the ground truth $PS$, and the 'best-fitting' $K^{trans}$ estimates had the highest accuracy compared to any of the models used alone. A smaller interstitial volume fraction led to worse underestimation of $K^{trans}$ by the Patlak model, and this effect was greater at high permeability. A smaller interstitial space will fill with contrast agent more quickly, leading to significant backflux, violating the key assumption of unidirectional contrast agent leakage described by the Patlak model and exacerbating the underestimation of $K^{trans}$.

The effect of variable $T_{10}$ on the accuracy of $K^{trans}$ estimation is shown in Supplementary Figure 5. In general, findings were comparable to the use of a fixed $T_{10}$ = 1000 ms representative of normal white matter, the selection of the preferred model was driven largely by the magnitude of the ground truth $PS$, and the 'best-fitting' $K^{trans}$ estimates had the highest accuracy compared to any of the models used alone. The Extended Tofts model overestimated $K^{trans}$ in the subtle permeability regime at lower $T_{10}$. It is possible that noise effects will be greater in a signal-time curve for a tissue with lower $T_{10}$, where $T_1$ shortening from the contrast agent results in smaller relative signal changes, possibly leading to a higher incidence of overfitting to noise at low permeability with the Extended Tofts model at lower $T_{10}$.

We next conducted Monte Carlo simulations, simultaneously varying all parameters with 1,000,000 sample combinations randomly selected within the specified physiological ranges, detailed in Supplementary Table 2. This is a more realistic representation of real DCE-MRI data, where underlying tissue will have varying ground truth parameter combinations in both health and disease. Supplementary Figure 6 shows binned averages of the ground truth vs fitted parameter relationships for $K^{trans}$, $v_p$, and $v_e$ across the model fitting approaches. In general, findings were comparable to the use of a fixed $F_p$, $v_p$, $v_e$, and $T_{10}$. The Patlak model underestimated $K^{trans}$ in the high permeability regime. The Extended Tofts model estimated $K^{trans}$ well in the high permeability regime, but had higher variability than the Patlak estimates at low permeability, while the model selection 'best-fitting' $K^{trans}$ estimates performed most accurately with lower relative variability. All models estimated $v_p$ similarly well, with the Patlak model marginally overestimating at high $v_p$. The Extended Tofts model estimates of $v_e$ were unreliable with very high variability, likely because $v_e$ is not readily measurable if permeability is low or noise is high, as is the case in these simulations. With the model selection approach, only concentration-time curves for sample combinations where the Patlak model is not appropriate (i.e. there is small $v_e$ and/or high *PS,* therefore backflux is detectable) are considered, reducing this variability.

**Supplementary Table 2.** Input values and physiological representations for Monte Carlo simulation simultaneously varying all parameters.

| Input parameter | Units | Distribution | Value Range (N samples) | Physiological Representation of Range |
|---|---|---|---|---|
| Permeability surface area product, *PS* | $min^{-1}$ | log-uniform | 0.0001 - 0.01 (1,000,000) | Subtle to very leaky BBB. |
| Plasma flow, $F_p$ | $min^{-1}$ | uniform | 0.1 – 1.2 (1,000,000) | Hypoperfused/ischaemic to highly perfused tissue. |
| Plasma volume fraction, $v_p$ | % | uniform | 1% – 10% (1,000,000) | Weakly vascularised to highly vascularised tissue. |
| Extravascular extracellular volume fraction, $v_e$ | % | uniform | 5% – 25% (1,000,000) | Small interstitial space (cell swelling/oedema) to large interstitial space (atrophy). |
| Baseline T1, $T_{10}$ | ms | uniform | 800 – 2000 (1,000,000) | Underlying brain tissue type (WM to blood). |

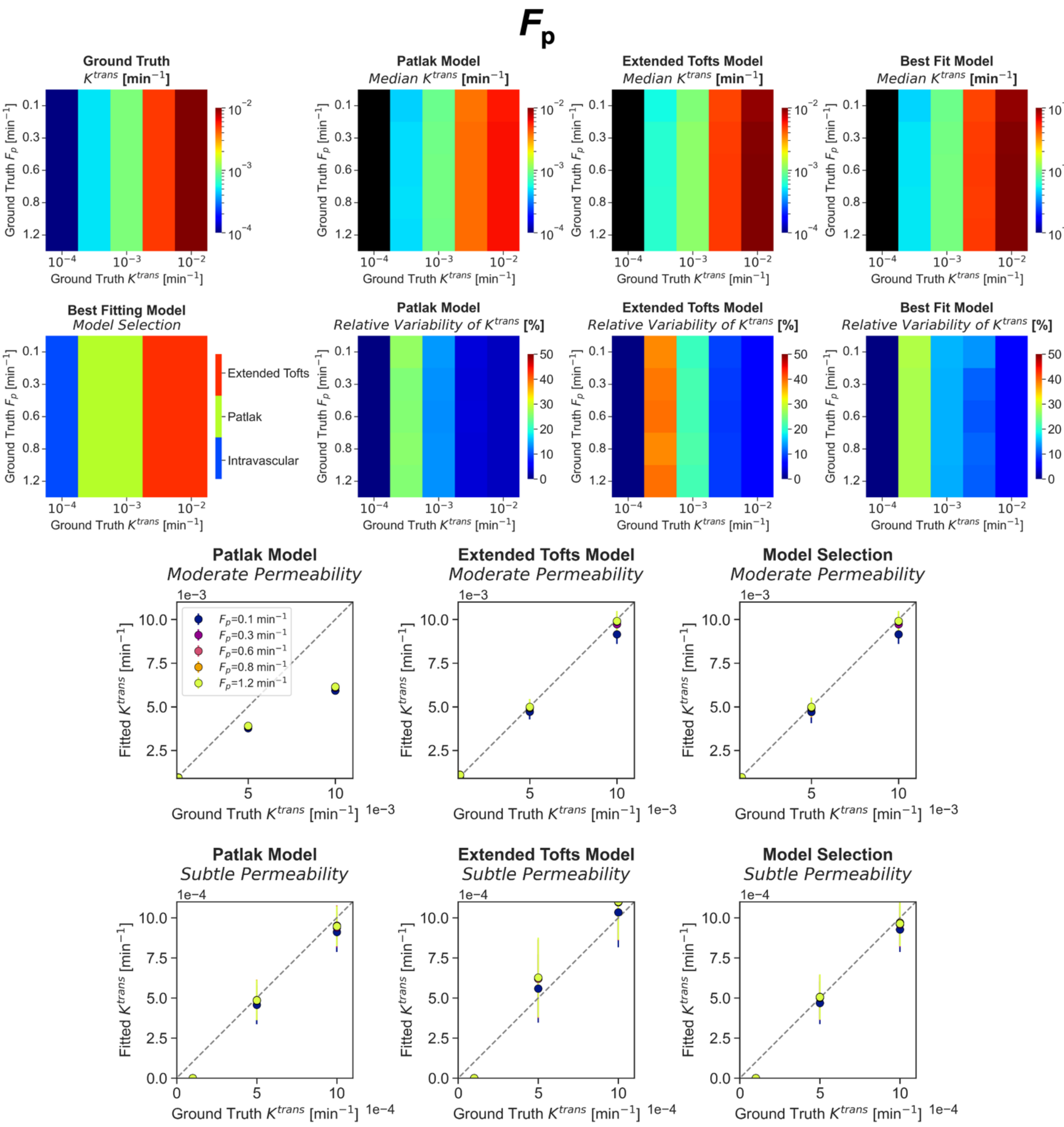


**Supplementary Figure 2. Model selection of noisy simulated data across a physiological range of $F_p$.** **(a)** Heatmap of the ground truth $K^{trans}$ input for the simulated concentration time curves. **(b)** Best-fitting model map for varying ground truth $K^{trans}$ and $F_p$. **(c – e)** For **(c)** the Patlak model, **(d)** the Extended Tofts Model, **(e)** Model Selection: **(i)** Heatmap of median fitted $K^{trans}$ estimates at varying ground truth $K^{trans}$ and $F_p$. **(ii)** Heatmap of relative variability (median absolute deviation as a percentage of the median) of $K^{trans}$ estimates at varying ground truth $K^{trans}$ and $F_p$. **(iii - iv)** For each $F_p$ input, median fitted $K^{trans}$ is plotted against the ground truth $K^{trans}$ in the (iii) high permeability and (iv) subtle permeability regimes. Error bars shown are the median absolute deviation.

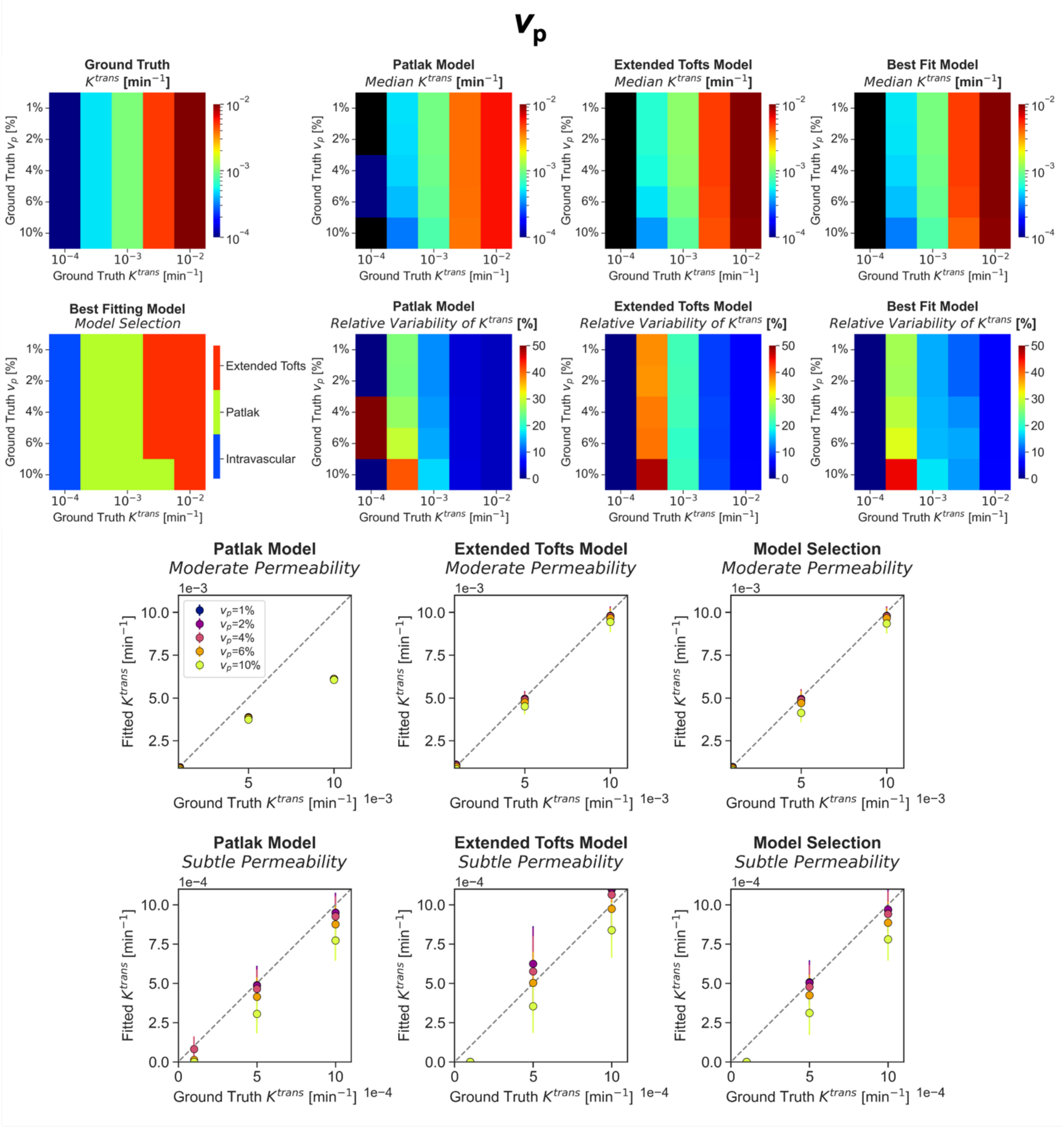


**Supplementary Figure 3. Model selection of noisy simulated data across a physiological range of $v_p$.** **(a)** Heatmap of the ground truth $K^{trans}$ input for the simulated concentration time curves. **(b)** Best-fitting model map for varying ground truth $K^{trans}$ and $v_p$. **(c – e)** For **(c)** the Patlak model, **(d)** the Extended Tofts Model, **(e)** Model Selection: **(i)** Heatmap of median fitted $K^{trans}$ estimates at varying ground truth $K^{trans}$ and $v_p$. **(ii)** Heatmap of relative variability (median absolute deviation as a percentage of the median) of $K^{trans}$ estimates at varying ground truth $K^{trans}$ and $v_p$. **(iii - iv)** For each $v_p$ input, median fitted $K^{trans}$ is plotted against the ground truth $K^{trans}$ in the (iii) high permeability and (iv) subtle permeability regimes. Error bars shown are the median absolute deviation.

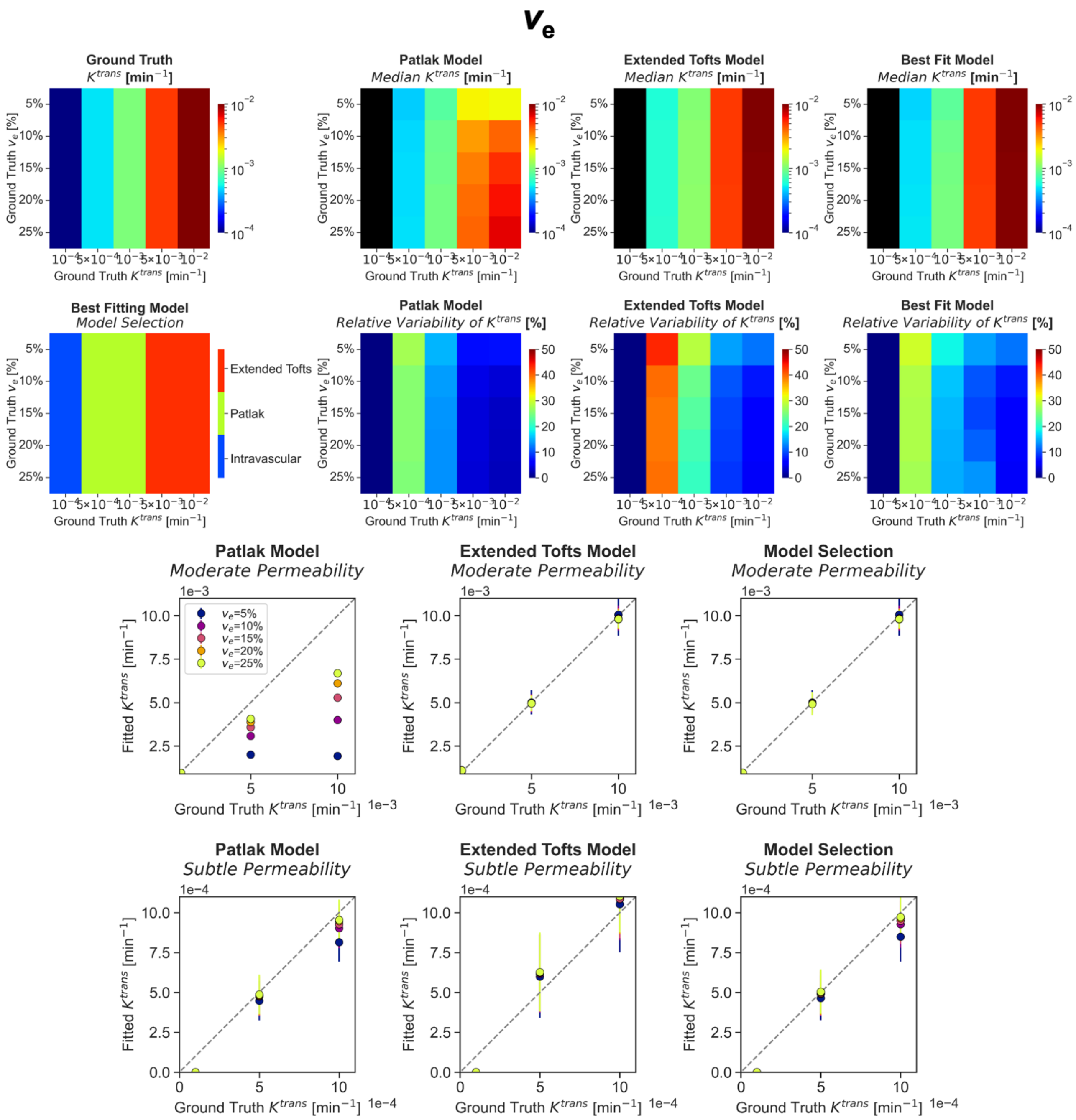


**Supplementary Figure 4. Model selection of noisy simulated data across a physiological range of $v_e$.** **(a)** Heatmap of the ground truth $K^{trans}$ input for the simulated concentration time curves. **(b)** Best-fitting model map for varying ground truth $K^{trans}$ and $v_e$. **(c – e)** For **(c)** the Patlak model, **(d)** the Extended Tofts Model, **(e)** Model Selection: **(i)** Heatmap of median fitted $K^{trans}$ estimates at varying ground truth $K^{trans}$ and $v_e$. **(ii)** Heatmap of relative variability (median absolute deviation as a percentage of the median) of $K^{trans}$ estimates at varying ground truth $K^{trans}$ and $v_e$. **(iii - iv)** For each $v_e$ input, median fitted $K^{trans}$ is plotted against the ground truth $K^{trans}$ in the (iii) high permeability and (iv) subtle permeability regimes. Error bars shown are the median absolute deviation.

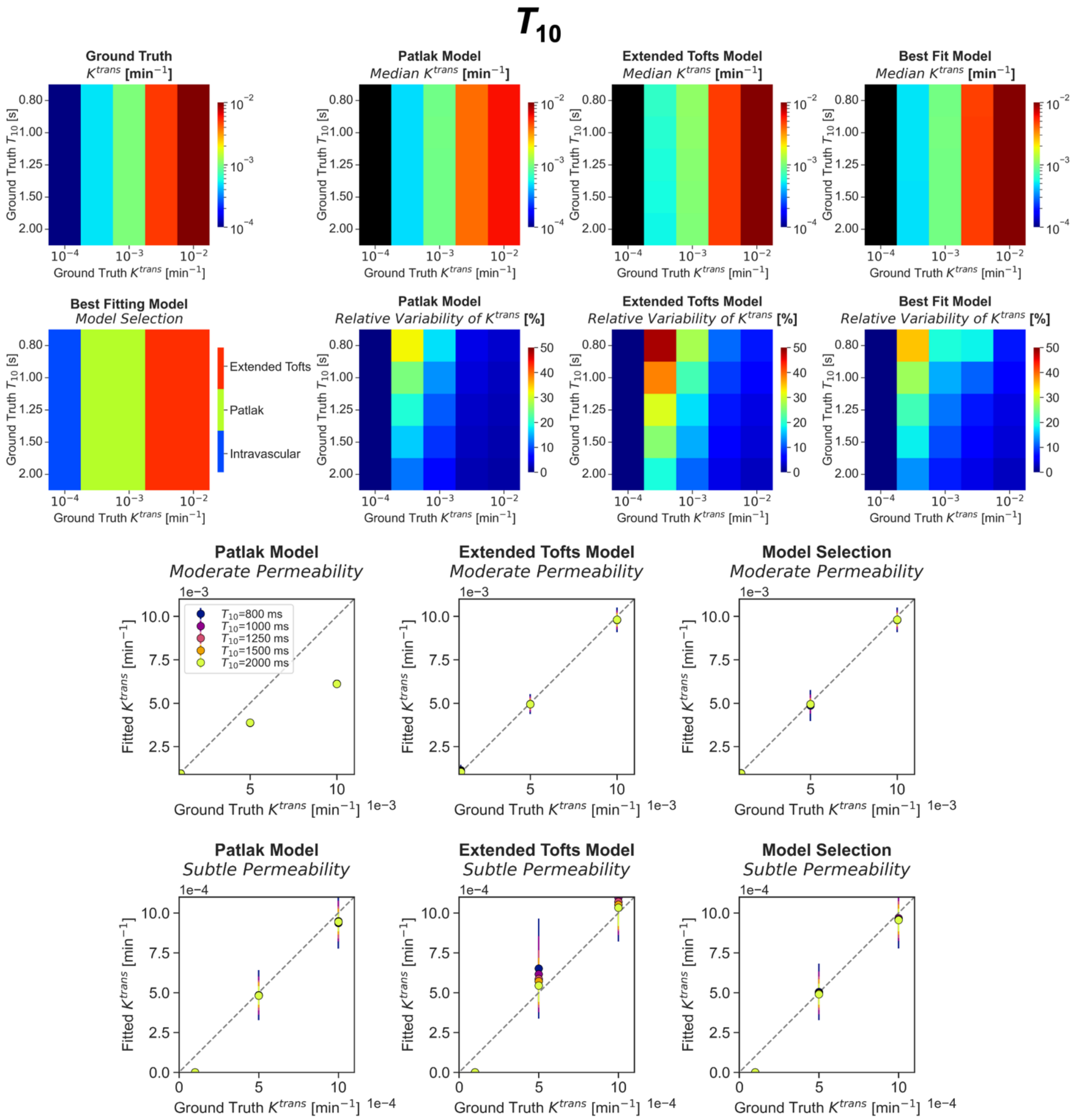


**Supplementary Figure 5. Model selection of noisy simulated data across a physiological range of $T_{10}$. (a)** Heatmap of the ground truth $K^{trans}$ input for the simulated concentration time curves. **(b)** Best-fitting model map for varying ground truth $K^{trans}$ and $T_{10}$. **(c – e)** For **(c)** the Patlak model, **(d)** the Extended Tofts Model, **(e)** Model Selection: **(i)** Heatmap of median fitted $K^{trans}$ estimates at varying ground truth $K^{trans}$ and $T_{10}$. **(ii)** Heatmap of relative variability (median absolute deviation as a percentage of the median) of $K^{trans}$ estimates at varying ground truth $K^{trans}$ and $T_{10}$. **(iii - iv)** For each $T_{10}$ input, median fitted $K^{trans}$ is plotted against the ground truth $K^{trans}$ in the (iii) high permeability and (iv) subtle permeability regimes. Error bars shown are the median absolute deviation.

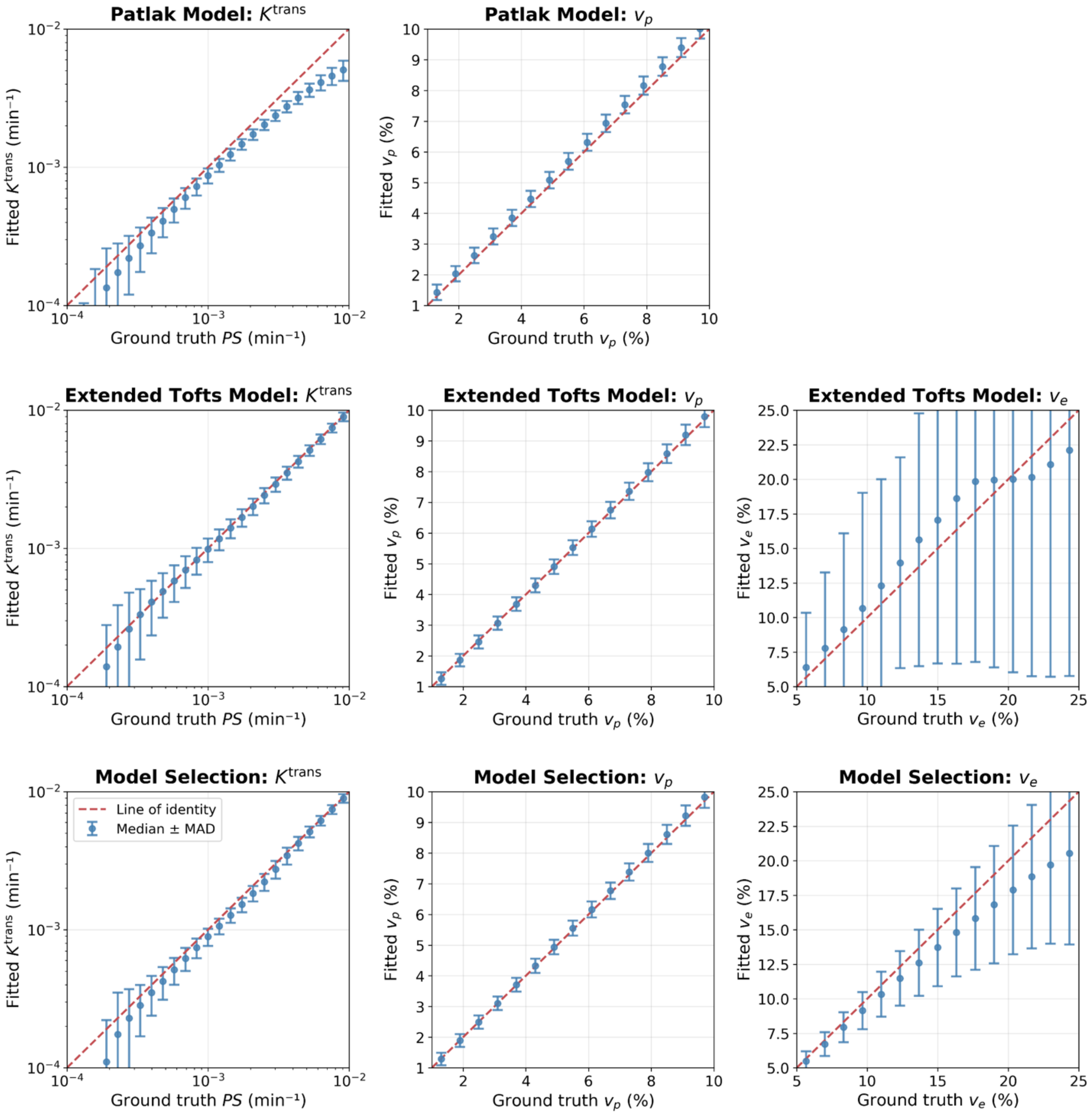


**Supplementary Figure 6. Model selection in Monte Carlo simulations varying all physiological input parameters. (a)** Patlak model estimates of $K^{trans}$ and $v_p$. **(b)** Extended Tofts model estimates of $K^{trans}$, $v_p$, and $v_e$. **(c)** Model selection estimates of ‘best-fitting’ $K^{trans}$, $v_p$, and $v_e$. Error bars are the median absolute deviation.
